# Psilocybin induces dose-dependent changes in functional network organization in rat cortex

**DOI:** 10.1101/2024.02.09.579718

**Authors:** Brian H. Silverstein, Nicholas Kolbman, Amanda Nelson, Tiecheng Liu, Peter Guzzo, Jim Gilligan, UnCheol Lee, George A. Mashour, Giancarlo Vanini, Dinesh Pal

**Author notes:** Equal Contribution (Giancarlo Vanini, Dinesh Pal). Correspondence: Dinesh Pal, Ph.D. 7433 Medical Science Building 1 1150 West Medical Center Drive Ann Arbor, Michigan 48109.

## Abstract

Psilocybin produces an altered state of consciousness in humans and is associated with complex spatiotemporal changes in brain networks. Given the emphasis on rodent models for mechanistic studies, there is a need for characterization of the effect of psilocybin on brain-wide network dynamics. Previous rodent studies of psychedelics, using electroencephalogram, have primarily been done with sparse electrode arrays that offered limited spatial resolution precluding network level analysis, and have been restricted to lower gamma frequencies. Therefore, in the study, we used electroencephalographic recordings from 27 sites (electrodes) across rat cortex (*n*=6 male, 6 female) to characterize the effect of psilocybin (0.1 mg/kg, 1 mg/kg, and 10 mg/kg delivered over an hour) on network organization as inferred through changes in node degree (index of network density) and connection strength (weighted phase-lag index). The removal of aperiodic component from the electroencephalogram localized the primary oscillatory changes to theta (4-10 Hz), medium gamma (70-110 Hz), and high gamma (110-150 Hz) bands, which were used for the network analysis. Additionally, we determined the concurrent changes in theta-gamma phase-amplitude coupling. We report that psilocybin, in a dose-dependent manner, 1) disrupted theta-gamma coupling [*p*<0.05], 2) increased frontal high gamma connectivity [*p*<0.05] and posterior theta connectivity [*p*≤0.049], and 3) increased frontal high gamma [*p*<0.05] and posterior theta [*p*≤0.046] network density. The medium gamma frontoparietal connectivity showed a nonlinear relationship with psilocybin dose. Our results suggest that high-frequency network organization, decoupled from local theta-phase, may be an important signature of psilocybin-induced non-ordinary state of consciousness.

## INTRODUCTION

Psilocybin is a serotonergic psychedelic that is being explored in clinical trials to treat psychiatric disorders.^1–6^ In humans, psilocybin produces an altered state of consciousness, associated with complex changes in brain network activity including increased posterior connectivity and decreased frontal connectivity,^7^ increased spatiotemporal complexity,^8–10^ decreased network segregation^11^ and decreased spectral power and connectivity in low (<40 Hz) frequencies.^12–14^ Functional connectivity changes have also been shown to correlate with psychedelic-induced subjective experiences, suggesting network dynamics may be an important component of the mechanisms underpinning the action of psychedelic drugs, including psilocybin.^12,15,16^ Recent electrophysiological studies in rodents suggest that the emergence of high gamma (>110 Hz) amplitude and connectivity, observed during administration of lysergic acid diethylamide (LSD), ketamine, and phencyclidine, may be an important common marker of psychedelics.^17–19^ However, this high gamma emergence phenomenon has not been documented during psilocybin administration and previous studies using psilocin or intracranial recordings after systemic delivery of psilocybin^20,21^ lacked the spatial resolution to allow corticocortical network analysis. Relatedly, gamma amplitude is known to be modulated according to the phase of theta band (4-10 Hz) oscillations, and it has been demonstrated that the relationship between these two frequency bands changes with cognitive demands and state of arousal.^22–25^ If local and interregional gamma-range dynamics are altered by psychedelic administration, disruption of theta-gamma coupling is likely a correlated phenomenon. However, psilocybin-induced theta-gamma decoupling has not been systematically demonstrated in rodents with co-occurring network-level changes.

Therefore, in this study, we used cortex-wide high-density (27 electrodes) electroencephalographic recordings to characterize the effect of psilocybin (0.1 mg/kg, 1 mg/kg, and 10 mg/kg) delivered intravenously over one hour, in Sprague Dawley rats (*n*=6 male and 6 female). We first performed spectral decomposition of the electroencephalogram (EEG) data to identify the primary oscillatory components that responded to psilocybin administration. These analyses showed that psilocybin altered three oscillatory components in the theta (4-10 Hz), medium gamma (70-110 Hz), and high gamma (110-150 Hz) bands, which were used for further analyzing the changes in phase-amplitude coupling and network organization. We report that psilocybin disrupted theta-gamma coupling and induced multiscale reorganization of cortical connectivity as shown by simultaneous increase in frontal high gamma connectivity and posterior theta connectivity, and a decrease in posterior medium gamma network connectivity. Our results also show increased posterior theta and frontal high gamma network density, which is a measure of the number of functional connections at any given electrode. These findings suggest that high-frequency network reorganization, decoupled from local theta-phase, may be an important signature of the non-ordinary state produced by psilocybin.

## RESULTS

All rats were implanted with electrodes to record EEG from across the cortex and received an hour-long intravenous infusion of three different doses of psilocybin and 0.9% saline (vehicle control), in a counterbalanced manner with at least 5-7 days of inter-experiment interval. We focused our analysis on 90 minutes of EEG data comprising a 10-minute pre-psilocybin infusion baseline period, 60 minutes spanning the intravenous psilocybin (0.1 mg/kg, 1 mg/kg, and 10 mg/kg) or 0.9% saline infusion time, and 20 minutes immediately after the end of psilocybin or saline infusion **(Figure 1)**.

**Figure 1.**
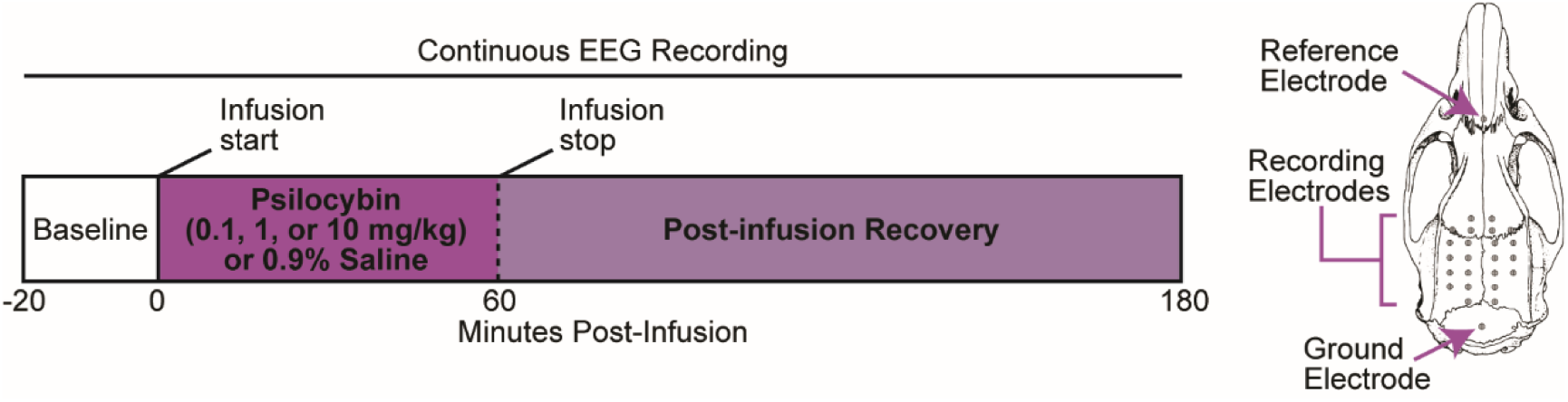
Schematic showing the experimental design and timeline. Each rat (*n*=12, 6 male, 6 female) received 0.9% saline and three doses of psilocybin (0.1 mg/kg, 1 mg/kg, and 10 mg/kg) as a continuous infusion over the course of an hour. Each infusion session was separated by 5-7 days and was conducted in a counter-balanced manner. On the right-side of the experiment timeline is shown an image of the rat cranium indicating the EEG, ground, and reference electrode location.

### Psilocybin altered spectral power in theta and gamma bands and dynamically shifted dominant EEG frequencies

Comparisons of the power spectrum between each dose of psilocybin and saline control indicated that psilocybin-related changes in spectral characteristics are not broadband but localized in frequency space and shifting over time (Figure 2A-C). In order to focus our analysis on the localized oscillatory changes, we applied the FOOOF algorithm^26^ to all EEG data. The FOOOF algorithm removes the aperiodic (1/f) component from the power spectrum, enabling unbiased estimation of the peak frequency and amplitude of oscillatory components (for representative spectral detrending see Figure 2D). This step is crucial to avoid confounding a change in the oscillatory peak frequency and amplitude with a change in the broadband component. Detrending the power spectrum revealed that the primary oscillatory components affected by psilocybin infusion corresponded to the theta (4-10 Hz), medium gamma (70-110 Hz), and high gamma (110-150 Hz) bands (Figure 2D).

**Figure 2.**
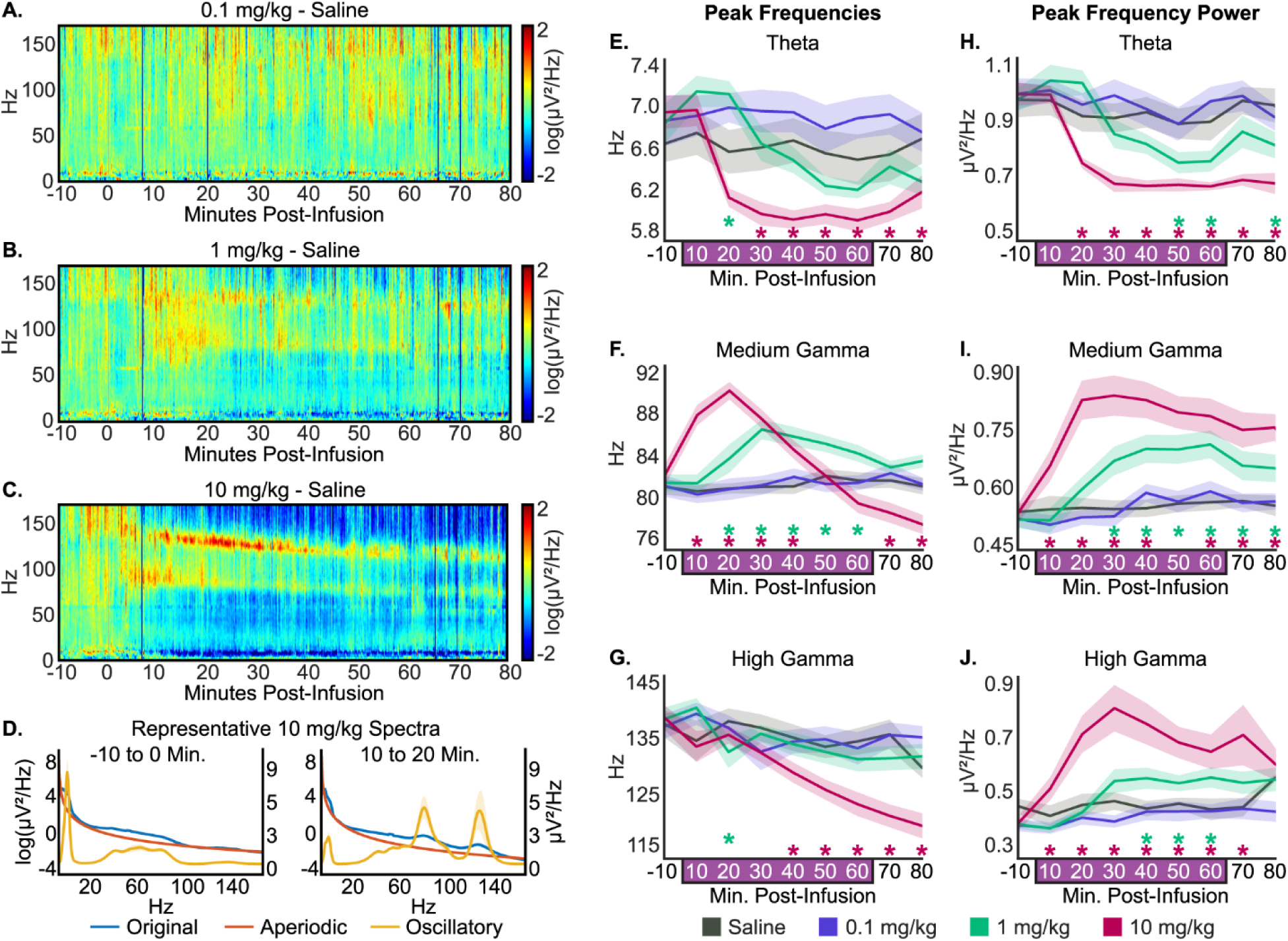
Psilocybin altered global peak oscillatory frequencies and amplitudes in a dose-dependent manner. (A-C) Global spectrograms averaged across rats (*n*=12, 6 male, 6 female) depicting the difference between each dose (0.1 mg/kg, 1 mg/kg, and 10 mg/kg) and saline. Psilocybin/saline infusion started at time=0 and stopped at time=60 minutes. Warm colors indicate higher spectral power while cool colors indicate lower spectral power relative to saline. (D) Representative global power spectrum averaged over minutes -10 to 0 (left) and 10-20 (right) of 10 mg/kg psilocybin dose. The FOOOF algorithm was used to model and remove the 1/f component from the original power spectrum to quantify band-specific peak frequencies and amplitudes. Displayed here are the original power spectrum (blue), the aperiodic component (red), and the oscillatory component (gold). For visualization the original and aperiodic spectra are log-transformed (left y-axis) and the oscillatory spectra are on a linear scale (right y-axis). (E-J) Changes in peak frequency and amplitude in the theta, medium gamma, and high gamma bands. The 1 mg/kg of psilocybin slowed theta peak frequency and increased medium gamma peak frequency. The 10 mg/kg psilocybin dose caused a significant decrease in theta peak frequency. The 10 mg/kg dose initially increased medium gamma peak frequency, but towards the end of infusion the peak frequency slowed relative to saline. The 1 mg/kg and 10 mg/kg doses of psilocybin decreased theta amplitude and increased medium and high gamma amplitudes relative to saline. The data are provided as mean ± standard error of the mean. \**p*<0.05, FDR-corrected post-hoc comparisons.

Analysis of the peak frequency and amplitude of each component over the duration of the recording period showed that there was no statistically significant effect of 0.1 mg/kg psilocybin infusion on EEG oscillatory components (*p*>0.05), but clear dose-dependent effects were observed during both 1 mg/kg and 10 mg/kg infusions (Figure 2E-J). Relative to saline infusion, peak frequency in the theta band (Figure 2E) briefly increased 10 minutes after the start of 1 mg/kg psilocybin infusion (*p*=0.0074), while the 10 mg/kg psilocybin produced a sustained decrease in theta peak frequency beginning after 30 minutes of infusion. The peak frequency in medium gamma band (Figure 2F) was significantly increased after 10 minutes of 1 mg/kg psilocybin infusion, which remained elevated (*p*≤0.039) for the duration of the infusion period. By contrast, the peak frequency in the medium gamma band (Figure 2F) showed a significant increase within the first 10 minutes of 10 mg/kg psilocybin infusion. The increase lasted for 40 minutes of infusion (*p*≤0.0014), after which the peak frequency decreased below saline levels (*p*≤0.0018). Finally, the high gamma peak frequency (Figure 2G) showed a transient decrease after 10 minutes of 1 mg/kg psilocybin infusion (*p=*0.036), and a sustained decrease after 30 minutes of 10 mg/kg psilocybin (*p*≤0.0026).

In addition to altering the peak oscillating frequency in theta, medium gamma, and high gamma bands, psilocybin infusion dose-dependently altered the peak amplitude in these bands. Relative to saline infusion, theta peak amplitude (Figure 2H) decreased after 40 minutes of 1 mg/kg psilocybin infusion (*p*≤0.020) and after only 10 minutes of 10 mg/kg psilocybin infusion (*p*≤0.039). In contrast, the peak amplitude in the medium gamma band (Figure 2I) showed a significant increase (*p*≤0.0032) after 20 minutes of 1 mg/kg psilocybin infusion and within 10 minutes of 10 mg/kg psilocybin (*p*≤0.00050), both of which remained elevated for the duration of the recording. Similarly, peak amplitude in the high gamma band (Figure 2J) showed a significant increase (*p*≤0.044) after 30 minutes of 1 mg/kg psilocybin infusion and within 10 minutes of 10 mg/kg infusion (*p*≤0.040), after which high gamma power remained elevated throughout the recording period (Figure 2J).

To reveal the relationship between changes in medium and high gamma power, and rat movement, we quantified rat activity – body or head movements – using a gyroscope built into the recording head-stage. Analysis of the mean 3D angular velocity throughout the recording session showed that rat movement was dissociated from the changes in theta and gamma-band power (Figure 3). As compared to saline infusion, 0.1 mg/kg psilocybin had no effect on rat movement (*p*>0.05). The 1 mg/kg dose induced a brief period (∼10 minutes) of increased movements after 10 minutes of infusion (*p*=0.0069), after which the level of movement returned to that observed after saline infusion. The 10 mg/kg psilocybin produced a significant decrease in rat movements between 30-60 minutes of infusion (all *p*≤0.023).

**Figure 3.**
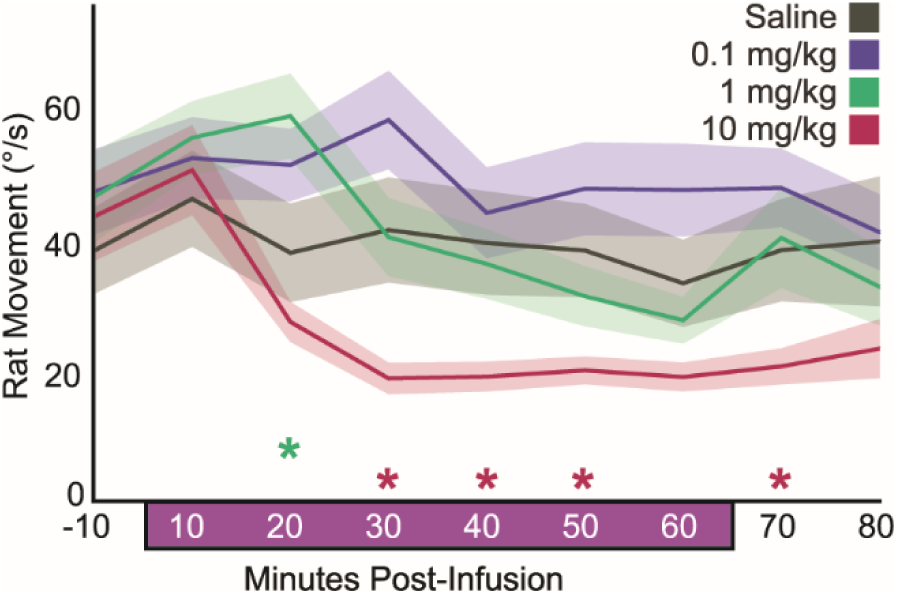
Effect of psilocybin on rat movement. 3D gyroscope activity averaged in 10-minute bins plotted as mean ± standard error of the mean. The 1 mg/kg dose briefly increased rat head movements, whereas the 10 mg/kg dose resulted in a quiescent state with minimal movement, thereby dissociating the increased gamma power from movement. \**p*<0.05, FDR-corrected post-hoc comparisons between saline and psilocybin doses.

In summary, spectral analysis of EEG dynamics during continuous psilocybin infusion indicated two distinct EEG states during the 10 mg/kg infusion. The initial period during 10 mg/kg infusion and later period during 1 mg/kg infusion were characterized by decreased theta and increased medium and high gamma power, along with a faster medium gamma peak frequency. As the 10 mg/kg infusion continued, it diverged from the lower dose, characterized by slowing theta and gamma peak frequencies and a notable increase in high gamma power. These data suggest that increasing doses of psilocybin may not have a linear effect on brain oscillatory dynamics.

### Psilocybin disrupted theta-gamma coupling in a dose-dependent manner

Theta- and gamma-range oscillations are known to be phase-amplitude coupled, with the phase of theta acting as a timing mechanism for local gamma amplitude.^22^ This coupling is believed to support cognitive functions, such as memory and attention, and is altered during changes in states of consciousness.^23–25^ Furthermore, recent studies have reported altered timing between spikes and the phase of local field potentials in rodents during lysergic acid diethylamide (LSD) and 2,5-Dimethoxy-4-Iodoamphetamine (DOI) administration. As compared to saline infusion, 0.1 mg/kg psilocybin did not produce any statistically significant changes in either theta-medium gamma or theta-high gamma PAC (*p*>0.05; Figure 4). In contrast, both 1 mg/kg and 10 mg/kg psilocybin infusion diminished theta-medium gamma PAC across the entire cortex (Figure 4A). After 20 minutes of 1 mg/kg infusion, there was a decrease in theta-medium gamma PAC in the extreme frontal (*p*≤0.047) and posterior/occipital areas (*p*≤0.049), which, during the last 20 minutes of 1 mg/kg infusion, spread to the rest of the cortex (*p*≤0.045). The PAC values started to recover back to saline values after the cessation of psilocybin infusion but remained significantly low at the end of 20 minutes post-psilocybin period (*p*≤0.038). The effect of 10 mg/kg psilocybin on theta-medium gamma PAC (Figure 4A) was similar, but the onset time was faster (10 minutes after 10 mg/kg vs 20 minutes after 1 mg/kg psilocybin). Interestingly, theta-high gamma PAC (Figure 4B) was only mildly affected by 1 mg/kg of psilocybin and with late onset (after 40 minutes of infusion). The decrease was restricted primarily to the frontal regions (*p*<0.050), which returned to saline values within 10 minutes after cessation of psilocybin infusion. The 10 mg/kg dose of psilocybin showed a much stronger effect on theta-high gamma PAC, which showed wide-spread decreases across the cortex after 10 minutes of infusion (*p*≤0.048) and which remained significantly low through the post-infusion period (Figure 4B). In summary, psilocybin infusion dose-dependently disrupted local theta-gamma coupling across the cortex, but particularly in frontal cortex, potentially dysregulating information coordination.

**Figure 4.**
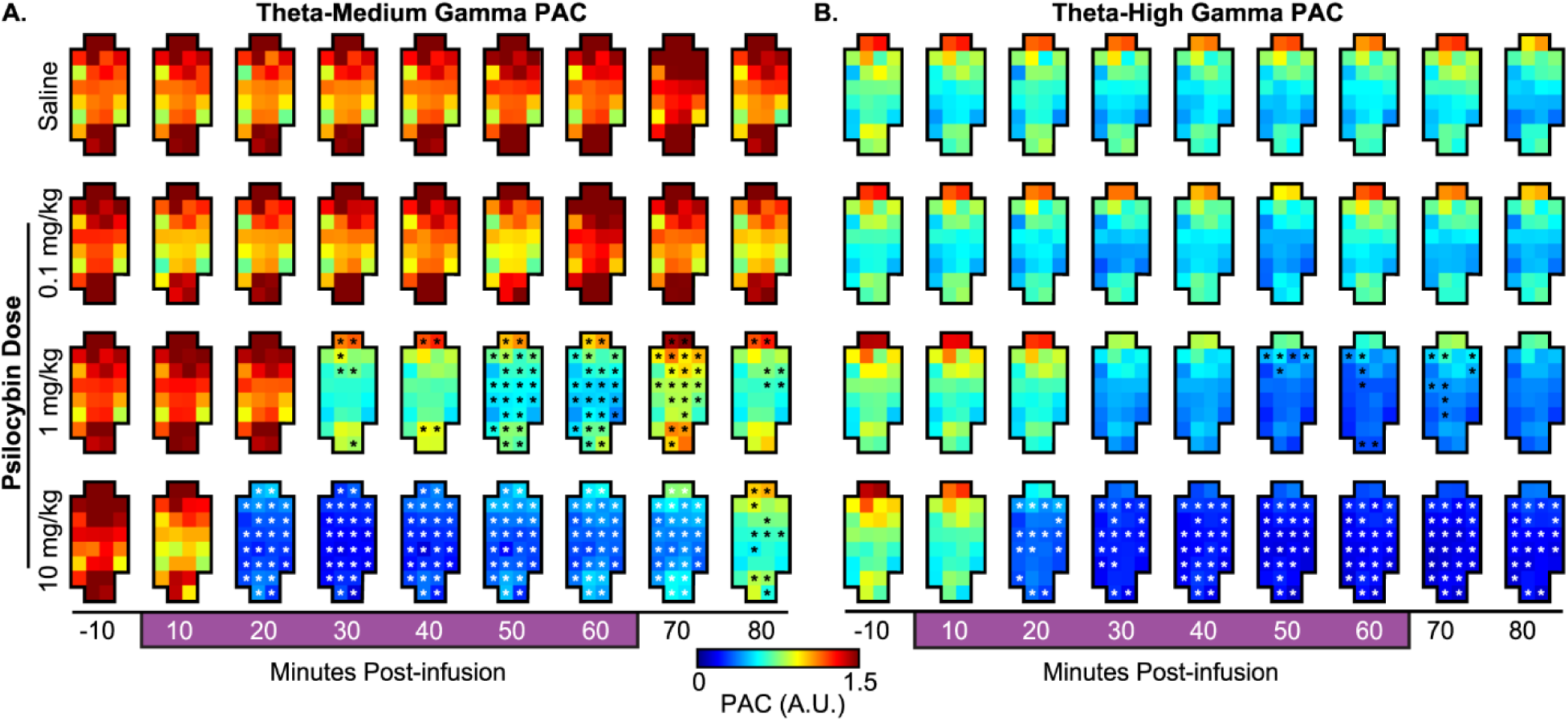
Intravenous psilocybin dose-dependently disrupted theta-gamma coupling. Following psilocybin infusion, both the 1 mg/kg and 10 mg/kg doses resulted in decoupling of theta phase from medium (A) and high (B) gamma amplitudes in a dose-dependent fashion. PAC was not altered by the 0.1 mg/kg dose. Each grid represents a 10-minute average of PAC values across all rats (*n*=12). Each square indicates an electrode in the layout described in Figure 1. White and black asterisks: *p*<0.05, FDR-corrected post-hoc comparisons.

### Psilocybin induced multi-scale reorganization of cortical connectivity in a dose-dependent manner

Functional network reorganization is a key component of current neuronal models of the psychedelic state.^27,28^ Therefore, next, we assessed if the psilocybin-induced dysregulation of local timing between theta and gamma oscillations is accompanied by reorganization of connectivity patterns and network structure in these frequency bands. To characterize psilocybin-induced changes in network organization, we assessed node degree, which is an index of network density or the number of functional connections at a given electrode, and edge-wise wPLI-based connection strength (Figure 5), before, during, and after psilocybin infusion.

**Figure 5.**
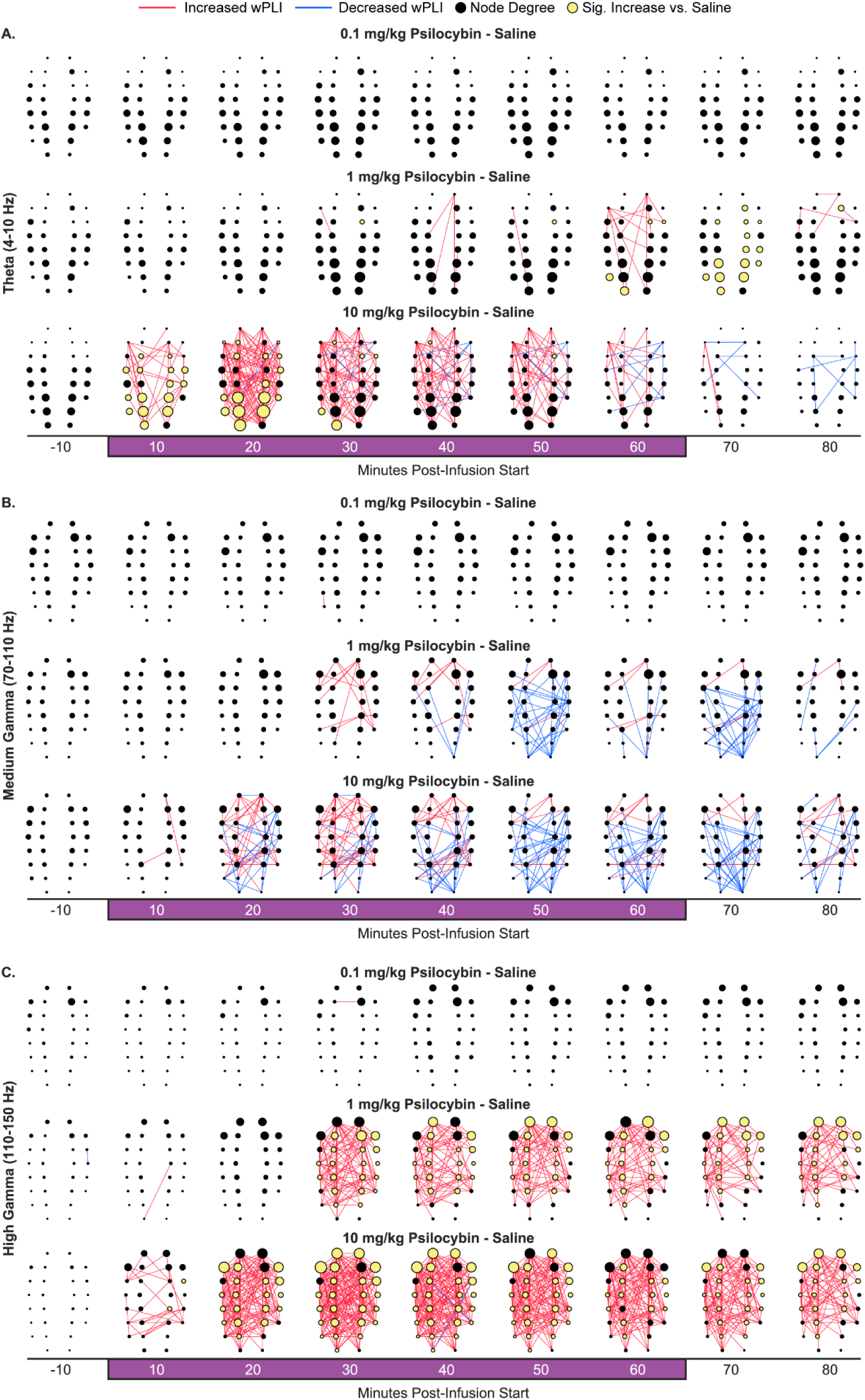
Intravenous psilocybin induced broad reorganization of theta and gamma cortical connectivity patterns. wPLI connectivity differences between psilocybin and saline averaged across rats (*n*=12) and over 10-minute bins. Red lines indicate increased wPLI relative to saline, blue lines indicate decreased wPLI. Dots indicate electrode location corresponding to the electrode map shown in Figure 1. Dot size indicates node degree magnitude. Yellow dots show significantly increased node degree relative to saline. Only connections that were significantly different (*p*<0.05) from saline following FDR-correction are displayed. Beginning halfway through infusion, the 1 mg/kg psilocybin dose caused sparse increases in frontoparietal theta connectivity (A), increased medium gamma frontal connectivity, but decreased posterior connectivity (B), and caused broad frontoparietal increases in high gamma connectivity (C). The 10 mg/kg psilocybin dose had a similar effect on cortical connectivity, but was amplified in a dose-dependent fashion, in particular causing large increases in theta posterior connectivity (A), increasing, then decreasing medium gamma frontoparietal connectivity (B), and increasing frontoparietal high gamma connectivity (C).

In accordance with the spectral and PAC analyses, there was no effect of 0.1 mg/kg psilocybin infusion on network density or network connectivity strength in any of the bands measured, while both 1 mg/kg and 10 mg/kg infusions resulted in pronounced multiscale network alterations (Figure 5). The posterior theta network density (Figure 5A) was transiently increased after 60 minutes of 1 mg/kg psilocybin infusion, before returning to saline levels within 20 minutes post-infusion (*p*≤0.045). By contrast, the 10 mg/kg psilocybin infusion rapidly increased theta network density, particularly at posterior electrodes, within 10 minutes of infusion (*p*≤0.046). This effect peaked after 20 minutes of infusion and returned to saline levels after 40 minutes of infusion. Paralleling the observed changes in theta network density, during the 1 mg/kg infusion, frontoparietal theta network connectivity was strengthened only across a few electrodes starting after 30 minutes infusion (*p*≤0.047), whereas the 10 mg/kg psilocybin infusion showed a strong, dynamic effect on theta frontoparietal networks. Theta connection strength increased within 10 minutes of 10 mg/kg infusion (*p*≤0.049), peaked after 30 minutes of infusion, then began to decrease, returning to saline levels by 20 minutes post-infusion.

Unlike the theta network, none of the psilocybin doses altered medium gamma network density relative to saline (*p*>0.050; Figure 5B). Although network density was constant, both the 1 mg/kg and 10 mg/kg doses of psilocybin had a biphasic effect on medium gamma frontoparietal network strength. Medium gamma frontal network synchronization was initially strengthened beginning after 20 and within 10 minutes of 1 mg/kg and 10 mg/kg infusions, respectively (*p*<0.050). After continued infusion, both the 1 mg/kg and 10 mg/kg doses of psilocybin induced widespread weakening of posterior-clustered medium gamma network connections beginning after 30 and 10 minutes of infusion, respectively (*p*<0.050).

High gamma network density and synchronization were strongly affected by psilocybin infusion (Figure 5C). Compared to saline, psilocybin caused a widespread increase in network density and synchronization strength, particularly in long-range connections to and from the frontal cortex, reflecting a marked change in brain network organization. Both the 1 mg/kg (*p*≤0.049) and 10 mg/kg (*p*<0.050) doses of psilocybin infusion resulted in increased frontal and posterior node degree beginning after 20 and 10 minutes of infusion, respectively. Similarly, local-and long-range frontoparietal synchronization was increased by both 1 mg/kg (*p*<0.050) and 10 mg/kg (*p*≤0.048) doses, again in a dose-dependent manner, beginning after 20 minutes and within 10 minutes of infusion, respectively.

In summary, psilocybin strengthened and broadened the frontal high gamma network in a dose-dependent manner, whereas psilocybin dose has a nonlinear effect on both posterior theta network density and connection strength, as well as medium gamma frontoparietal connectivity, suggesting two distinct states for lower and higher doses of psilocybin. These reorganized connectivity patterns may be enabled by psilocybin-associated loss of local PAC-based modulation of theta and gamma band temporal dynamics.

## DISCUSSION

Large-scale network reorganization is consistently reported in human psychedelic studies,^8,10,11,29^ and although the particular dynamics reported can vary considerably, reconfigured cortical network dynamics are believed to play a role in enabling the psychedelic-associated altered state of consciousness. Yet, without corresponding rodent models of psychedelic-related network dynamics, establishing mechanistic targets is a challenge. In this study, we report that intravenous psilocybin infusion has a nonlinear effect on brain network dynamics, resulting in two distinct brain network configurations that emerged over the course of infusion. The 1 mg/kg infusion and the first half of the 10 mg/kg infusion period were primarily characterized by a high-density posterior theta and frontal high gamma network, with strengthened frontoparietal medium gamma connectivity, and cortex-wide decoupling of theta phase from gamma amplitudes. During the later portion of the 10 mg/kg infusion, the brain network state became dominated by the high gamma network as the posterior theta density decreased and frontoparietal medium gamma connectivity continued to weaken.

To our knowledge, this is the first study to report network dynamics during psilocybin administration in rodents. In a recent study, Vejmola and colleagues^20^ reported cortex-wide decease in spectral power below 40 Hz and localized decreases in corticocortical coherence below 14 Hz after subcutaneous psilocin – the active metabolite of psilocybin – administration in rats.^20^ However, this study used a sparse electrode array with limited spatial resolution, which precluded network analysis, and was limited to low frequency (<40 Hz) oscillations. Of note, the divergence in theta band connectivity results between this study and ours is likely due to methodological differences. Instead of coherence, our study utilized wPLI, which is robust to volume conduction and therefore may differ from coherence especially in low-frequency, high-amplitude bands such as theta. Furthermore, differences in brain dynamics may be due to the different routes of administration or the use of psilocin versus psilocybin.

A notable feature of the network results presented here is the simultaneous change in the theta posterior network and the frontal high gamma network, which partially overlap. Several reports of human fMRI data indicate that psilocybin likely results in a reorganization of functional network topology.^8,30^ The increased frontal high gamma network density, nonlinear changes in posterior theta network density, and broad remodulation of frontoposterior connectivity strengths across theta and gamma bands are consistent with human fMRI data indicating increased long-range between-network connectivity,^11^ as well as reports that during psilocybin administration the spatial distribution of long-range networks varies over time.^10,16^

A growing body of literature emphasizes the role of frontoposterior high gamma connectivity during psychedelic administration in rodents and humans.^14,17,21,28,31^ Increased high gamma frontoposterior and frontostriatal connectivity has been reported in rodents administered serotonergic psychedelics such as LSD, DOI, and atypical psychedelics such as ketamine, nitrous oxide, and phencyclidine.^17–19,32^ These long-range networks may play an important role in facilitating a psychedelic state; for example, in humans, increased long-range connectivity is correlated with blood plasma levels of psilocybin and subjective drug experience.^11,12^

Local cortical circuit conditions also play a significant role in determining large-scale network synchronization. A few rodent studies of 5HT2A- and NMDA-associated psychedelics have used depth electrodes to localize high gamma activity to sites in frontal cortex, including olfactory cortex, orbital cortex, medial prefrontal cortex, and anterior cingulate cortex, as well as in parietal cortical areas of sensorimotor cortex and subcortical sites in striatum and nucleus accumbens.^17,21,32^ Similarly, decoupling of local spiking activity from the phase of low frequency local field potentials has also been observed following psilocybin, LSD, DOI, ketamine, and phencyclidine administration in rodents,^17,21^ which parallels the dose-dependent loss of theta-gamma coupling observed in our data. Gamma peak frequency may be partially determined by the balance of excitatory and inhibitory activity^33^ and has been shown to be an important index of changing cortical conditions associated with both cognitive demand and level of arousal.^34^ In humans, psilocybin has been reported to increase GABA and glutamate concentrations in the hippocampus and decrease both in the medial prefrontal cortex.^31^ In rodents, LSD and DOI are demonstrated to down-modulate both interneurons and pyramidal cell spiking activity.^17^

Finally, the increased high gamma network activation and associated increased gamma power may also have relevance for the neuroplastic effects of psilocybin. Recent evidence has indicated that psychedelics activate BDNF and mTOR signaling pathways which result in structural changes such as acute and lasting increases in dendritic spine density.^35–37^ Synaptic plasticity has also been associated with the same increased spiking activity^38^ that generates local increases in gamma (>70 Hz) power.^39,40^ However, further research is necessary to directly link psychedelic-induced high gamma activity to changes in neuroplasticity.

There are several caveats to our study. First, the increase in connectivity between association cortices does not rule out significant roles for the claustrum, basal ganglia, thalamus, or cerebellum, which are key to some models of psychedelic action.^27,41^ Second, because wPLI does not provide direction of influence, our data do not provide evidence for a “top down” or “bottom up” mechanism for the psilocybin state. Lastly, it is important to note that due to the lack of verbal report or an objective measure to report ‘content’ of consciousness in animals, we cannot comment on the presence or absence of psychedelic experience.

Overall, our results demonstrate nonlinear, multiscale changes in network organization, which are consistent with fMRI reports of shifting patterns of decreased within-network and increased between-network connectivity.^11,15,30^ The two distinct phases of theta network density and medium gamma network strength, suggest that rather than having a linear effect on brain network dynamics, higher doses of psilocybin likely produce a brain state distinct from lower doses.

## METHODS

Adult Sprague Dawley rats (*n*=12, 6 female, 6 male, weight 300-350g, Charles River Laboratories Inc., Wilmington, MA) maintained on 12:12 light: dark cycle (lights on at 8:00 am) and with *ad libitum* food and water, were used for all experiments. The experiments were approved by the Institutional Animal Care and Use Committee at the University of Michigan, Ann Arbor, and were conducted in compliance with the Guide for the Care and Use of Laboratory Animals (Ed 8, National Academies Press) and ARRIVE Guidelines.^42^

### Surgical procedures

Under surgical isoflurane (Piramal Enterprises, Telangana, India) anesthesia, rats were implanted with stainless steel screw electrodes (J.I. Morris Miniature Fasteners, Oxford, MA, USA; #FF000CE094) to record electroencephalogram (EEG) from 27 sites across the cortex and bilateral wire electrodes (Cooner Wire Company, Chatsworth, CA, USA; # AS 636) to record electromyogram (EMG) from dorsal nuchal muscles. An in-dwelling chronic catheter (Micro-Renathane tubing, MRE-040, Braintree Scientific, MA) was positioned in the jugular vein for infusion of psilocybin (Cayman Chemical, MI; CAS 520-52-5) and 0.9 % saline (Hospira, Lake Forest, IL, USA; #00409-4888-20). The jugular venous catheter was flushed with 0.2 mL of heparinized (1 unit/mL, Sagent Pharmaceuticals, Schaumburg, IL) saline and locked with 0.05 mL of Taurolidine-Citrate Catheter lock solution (TCS-04, Access Technologies, Skokie, IL) every 5-7 days to maintain catheter patency. Subcutaneous buprenorphine hydrochloride (Buprenex ®, Reckitt Benckiser Pharmaceuticals Inc., Richmond, VA; 0.01 mg/kg) and carprofen (Hospira, Inc., Lake Forest, IL; 5 mg/kg) were used for pre-surgical analgesia. Post-surgical analgesia was achieved with buprenorphine hydrochloride (0.03 mg/kg) administration every 6-8 h for 48 h. All rats were allowed at least 7-10 days of post-surgical recovery, during which they were conditioned to the EEG acquisition system.

### Experimental design

The experimental design is illustrated in Figure 1. All experiments were conducted between 9:00 am and 2:00 pm. On the day of the experiment, rats were connected to the EEG recording system and allowed to habituate for a minimum of 1 h. Following habituation, baseline EEG was acquired for 20 minutes. Thereafter, one of the three doses of psilocybin (0.1, 1.0, or 10 mg/kg) was delivered (intravenous) over an hour. On a separate day, the rats received 0.9% saline (vehicle control) for over an hour. The EEG data were collected continuously throughout the baseline, psilocybin/saline delivery, and for 60 minutes post-infusion. To maintain a constant behavioral state, rats were kept awake throughout the procedure with gentle tapping on the recording chamber. All rats received each of the three doses of psilocybin as well as vehicle control (0.9% saline), in a counter-balanced manner with a minimum of 5-7 days in between experiments to allow for drug washout.

### Electrophysiological data acquisition

Electrophysiological signals were recorded using the Cereplex Direct recording system (Blackrock Neurotech, Salt Lake City, UT). Monopolar EEG (0.1-300 Hz, sampling rate 1 kHz) and bipolar EMG (bandpass filtered between 0.1 and 125 Hz and sampled at 500 Hz) were recorded continuously throughout the experiment. The recording head-stage was equipped with a motion sensor for which the data were bandpass filtered between 0.1-50 Hz and sampled at 500 Hz. EEG data were manually inspected for quality; segments or channels with excessive noise were excluded from further analysis. Finally, EEG data were downsampled to 500 Hz prior to computational analysis.

### Spectral analysis

Spectral characteristics of the EEG data were estimated at each channel in 2-second non-overlapping windows using a multitaper wavelet decomposition as implemented in Fieldtrip.^43^ Using the FOOOF algorithm,^26^ we estimated the aperiodic offset and aperiodic exponent of the power spectrum and removed the 1/f component to facilitate peak frequency detection. Maximum peaks were estimated for the theta (4-10 Hz), medium gamma (70-110 Hz), and high gamma (110-150 Hz) bands. For statistical analysis, frequency band amplitude and peak frequency were averaged across 10-minute non-overlapping windows and across channels for global analysis.

### Phase-amplitude coupling

Phase-amplitude coupling (PAC) was estimated for artifact-free 10-second epochs according to the method first described by Canolty,^44^ and expanded by Onslow.^45^ PAC was computed between low frequencies (*LF*), bandpass filtered in 2 Hz steps, centered from 2 to 18 Hz, and high frequencies (*HF*), bandpass filtered in 5 Hz steps, centered from 27 to 197 Hz. Data were filtered and converted into analytic signals using a wavelet convolution (Morlet wavelet, width=7). The instantaneous phase, *θ*, and amplitude, *A*, were extracted from each signal and combined to create a third composite signal, *X*, such that 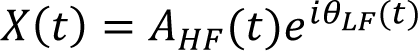 where *t* is time. In a given window and channel, PAC was calculated as 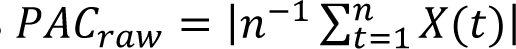 where *n* is the number of samples in a window. To ensure that values of PAC are independent of large power fluctuations or nonuniform phase distributions, we then applied permutation testing to generate normalized values of PAC in each window and channel. In each window, the amplitude time series extracted from the Morlet wavelet transformation were shuffled prior to computing a new PAC value, *PAC_shuff_*. This procedure was repeated 50 times, then the mean, σ_*shuff*_, were used to compute normalized PAC such that 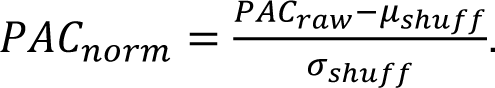. Mean PAC_norm_ was computed for each 10-minute epoch at each channel and at the global level. PAC_norm_ values in specific frequency band pairs, e.g. theta-medium gamma coupling, were computed by averaging PAC_norm_ values within each frequency-frequency range.

### Weighted phase-lag index

Weighted phase-lag index (wPLI) is a measure of functional connectivity that estimates the consistency of the phase relationship between two signals and is robust to volume conduction.^46^ To compute wPLI for a given channel pair of electrodes 𝑥 and 𝑦, we first applied a FIR bandpass filter and Hilbert transform to the monopolar EEG data to extract the analytic signal of each electrode. The complex conjugate of each channel pair was next computed to estimate the cross spectrum, *C_xy_*. Next, taking the imaginary component of 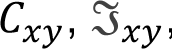, wPLI was then estimated as 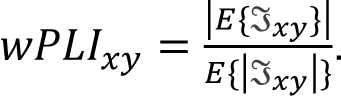. To rule out spurious connectivity due to the spectral distribution, we shuffled the phases of each channel while maintaining the amplitude distribution, using an FFT-based approach,^47^ to compute *wPLI_shuff_*. Analogous to the approach taken with PAC, this procedure was repeated 50 times, then the mean, 𝜇_*shuff*_, and standard deviation, 𝜎_*shuff*_, were used to compute normalized wPLI such that 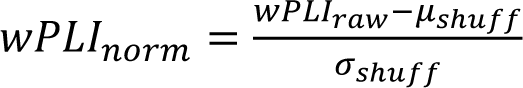. The channel-pair values of *wPLI_norm_* were averaged to estimate global *wPLI_norm_*. Local and global estimates of *wPLI_norm_* were averaged within each 10-minute epoch for statistical analysis. Finally, to compute node degree, binary undirected networks were computed from wPLI_norm_ by binarizing each 27 x 27 matrix according to a threshold of *p*=0.05, equivalent to wPLI_norm_=1.67. Node degree is then computed by taking the sum of each row.

### Statistical analysis

Statistical analyses were conducted in consultation with *the Consulting for Statistics, Computing and Analytics Research* unit at the University of Michigan (Ann Arbor, Michigan) using R (version 4.0.2). To facilitate statistical analysis, each EEG measure was averaged into 10-minute non-overlapping windows. Linear mixed models were computed at the global level for spectral analysis with rat as a random factor and i) dose, ii) time, iii) weight, and iv) sex as fixed factors. For channel-level analysis, the model was repeated for each channel, post-hoc comparisons between saline and each drug condition were extracted, and associated *p*-values were false discovery rate (FDR)-adjusted with an alpha=0.05.

## Acknowledgements

We thank Dr. Chris Andrews (Consulting for Statistics, Computing & Analytics Research, University of Michigan, Ann Arbor, Michigan) for consultation and help with the statistical analysis.

## Author Contributions

B.H.S., G.A.M., G.V., and D.P. designed the research; N.K and T.L. performed the experiments; B.H.S. and A.N. analyzed the data; all authors contributed to drafting the manuscript; B.H.S., U.L., G.A.M., G.V., and D.P. interpreted the data and wrote the manuscript.

## Data Availability Statement

All data and code used to analyze the data and conduct statistical comparisons are available from the authors upon request.

Declaration of interest: Jim Gilligan, Ph.D. is President & Chief Scientific Officer at Tryp Therapeutics. Peter Guzzo, PhD is Consulting VP, Drug Development at Tryp Therapeutics.

## Funding

This research was supported by the National Institutes of Health Grant R01 GM111293 to G.A.M. and D.P., funding from Tryp Therapeutics, San Diego, CA to G.V. and D.P., and the Department of Anesthesiology, University of Michigan Medical School, Ann Arbor, MI.

## Notes

### Summary of Updates

Correcting a figure legend typo.

